# Brain computation by assemblies of neurons

**DOI:** 10.1101/869156

**Authors:** Christos H. Papadimitriou, Santosh S. Vempala, Daniel Mitropolsky, Michael Collins, Wolfgang Maass

## Abstract

Assemblies are large populations of neurons believed to imprint memories, concepts, words and other cognitive information. We identify *a repertoire of operations* on assemblies. These operations correspond to properties of assemblies observed in experiments, and can be shown, analytically and through simulations, to be realizable by generic, randomly connected populations of neurons with Hebbian plasticity and inhibition. Operations on assemblies include: projection (duplicating an assembly by creating a new assembly in a downstream brain area); reciprocal projection (a variant of projection also entailing synaptic connectivity from the newly created assembly to the original one); association (increasing the overlap of two assemblies in the same brain area to reflect cooccurrence or similarity of the corresponding concepts); merge (creating a new assembly with ample synaptic connectivity to and from two existing ones); and pattern-completion (firing of an assembly, with some probability, in response to the firing of some but not all of its neurons). Our analytical results establishing the plausibility of these operations are proved in a simplified mathematical model of cortex: a finite set of brain areas each containing *n* excitatory neurons, with *random* connectivity that is both recurrent (within an area) and afferent (between areas). Within one area and at any time, only *k* of the *n* neurons fire — an assumption that models inhibition and serves to define both assemblies and areas — while synaptic weights are modified by Hebbian plasticity, as well as homeostasis. Importantly, all neural apparatus needed for the functionality of the assembly operations is created on the fly through the randomness of the synaptic network, the selection of the *k* neurons with the highest synaptic input, and Hebbian plasticity, without any special neural circuits assumed to be in place. Assemblies and their operations constitute a computational model of the brain which we call the *Assembly Calculus*, occupying a level of detail intermediate between the level of spiking neurons and synapses, and that of the whole brain. As with high-level programming languages, a computation in the Assembly Calculus (that is, a coherent sequence of assembly operations accomplishing a task) can ultimately be reduced — “compiled down” — to computation by neurons and synapses; however, it would be far more cumbersome and opaque to represent the same computation that way. The resulting computational system can be shown, under assumptions, to be in principle capable of carrying out arbitrary computations. We hypothesize that something like it may underlie higher human cognitive functions such as reasoning, planning, and language. In particular, we propose a plausible brain architecture based on assemblies for implementing the syntactic processing of language in cortex, which is consistent with recent experimental results.

## 1 Introduction

How does the brain beget the mind? How do molecules, cells, and synapses effect cognition, behavior, intelligence, reasoning, language? The remarkable and accelerating progress in neuroscience, both experimental and theoretical-computational, does not seem to bring us closer to an answer: the gap is formidable, and seems to necessitate the development of new conceptual frameworks. As Richard Axel recently put it [1] *“we do not have a logic for the transformation of neural activity into thought and action. I view discerning [this] logic as the most important future direction of neuroscience”*.

What kind of formal system, embodying and abstracting the realities of neural activity, would qualify as the sought “logic”?

We propose a formal computational model of the brain based on *assemblies of neurons;* we call this system the *Assembly Calculus*. In terms of detail and granularity, the Assembly Calculus occupies a position intermediate between the level of individual neurons and synapses, and the level of the whole brain models useful in cognitive science, e.g. [30, 31].

The basic elementary object of our system is the *assembly of excitatory neurons*. The idea of assemblies is, of course, not new. They were first hypothesized seven decades ago by Donald O. Hebb [20] to be densely interconnected sets of neurons whose loosely synchronized firing in a pattern is coterminous with the subject thinking of a particular concept or idea. Assembly-like formations have been sought by researchers during the decades following Hebb’s prediction, see for example [2], until they were clearly identified more than a decade ago through calcium imaging [17, 5]. More recently, assemblies (sometimes called *ensembles*) and their dynamic behavior have been studied extensively in the animal brain, see for example [27].

Our calculus outfits assemblies with certain *operations* that create new assemblies and/or modify existing ones: *project, reciprocal-project, associate, merge*, and a few others. These operations reflect known properties of assemblies observed in experiments, and they can be shown, either analytically or through simulations (more often both), to result from the activity of neurons and synapses. In other words, the high-level operations of this system can be “compiled down” to the world of neurons and synapses — a fact reminiscent of the way in which high-level programming languages are translated into machine code.

### Model

Our mathematical results, as well as most of our simulation results, employ a simplified and analytically tractable yet plausible model of neurons and synapses. We assume a finite number of brain areas denoted *A, B, C*, etc., intended to stand for an anatomically and functionally meaningful partition of the cortex (see Figure 2). Each area contains a population of *n* excitatory neurons^1^ with *random* recurrent connections. By this we mean that each ordered pair of neurons in an area has the same small probability *p* of being connected by a synapse, independently of what happens to other pairs — this is a well studied framework usually referred to as *Erdős–Renyi graph* or *G*_*n,p*_ [11]. In addition, for certain ordered pairs of areas, say (*A, B*), there are random *afferent* synaptic interconnections from *A* to *B*; that is, for every neuron in *A* and every neuron in *B* there is a chance *p* that they are connected by a synapse.^2^

**Figure 1:**
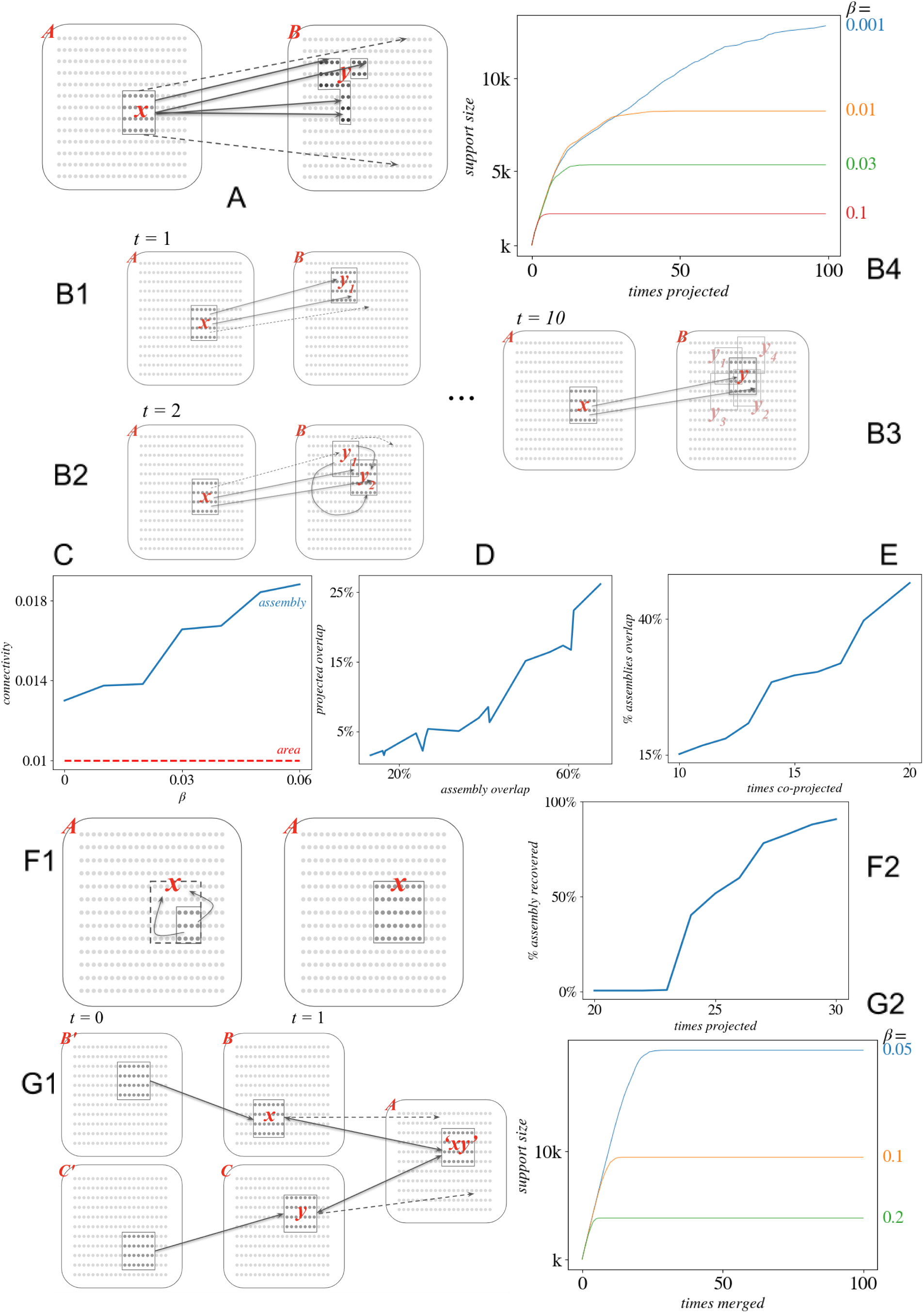
Assembly operations. **A RP&C:** If an assembly *x* fires in area *A*, and there is afferent connectivity to area *B*, the ensuing synaptic input will cause a set of *k* neurons in area *B*, call it *y*, to fire. The set *y* contains the *k* neurons in area *B* that receive the highest synaptic input from assembly *x*. This is an important primitive called *random projection and cap* (RP&C). **B1 – B3:** If assembly *x* fires again, afferent synaptic input from area *A* will be combined with recurrent synaptic input from *y*_1_ in area *B* to cause a new set of *k* neurons in area *B, y*_2_, to fire. Continued firing of *x* will create a sequence of sets of firing neurons in *B*: *y*_1_, *y*_2_, *y*_3_, … In the presence of Hebbian plasticity, this sequence converges exponentially fast to a set of neurons, an assembly *y* which we call the *projection of x to B*. If subsequently *x* fires, *y* will follow suit. **B4 Exponential convergence of assembly projection:** The horizontal axis is the number of times assembly *x* fires; the vertical axis is the total number of neurons in area B that fired in the process. Different color lines represent different values of the plasticity coefficient *β*; for higher levels of *β*, convergence is faster. **C**: Synaptic density within the resulting assembly increases with higher values of *β* (blue line), and is always higher than the baseline random synaptic connectivity *p* (dotted red line). **D Preservation of overlap:** Assembly overlap is preserved under RC&P. The x axis represents the overlap in two assemblies that are then projected to a new area once (RC&P); the y axis represents the percentage overlap of the two resulting assemblies in B. **E Association:** If two assemblies in two different areas have been independently projected in a third area to form assemblies *x* and *y*, and subsequently the two parent assemblies fire simultaneously, then each of *x, y* will respond by having some of its neurons migrate to the other assembly; this is called *association* of *x* and *y*. Such overlap due to association may reach 8 *-* 10% [21]. The figure shows results from simulation: first, two separate, stable assemblies are created in two areas A and B. Then, the assemblies in A and B are projected simultaneously into C for x time steps, represented by the x axis; the y axis shows the resulting overlap in the projected assemblies of A and B after the simultaneous firing. **F1 Pattern completion:** The firing of a few cells of an assembly results in the firing of the whole assembly. **F2:** An assembly is created in an area A by repeated firing of its parent; the number of times the parent fires is depicted in the horizontal axis. Next, 40% of neurons of the assembly are selected at random and fire for a number of steps: the y axis shows the overlap of the resulting assembly with the original assembly. With more reinforcement of the original assembly, the subset recovers nearly all of the original assembly. **G1 Merge:** The most advanced, and demanding in its implementation, operation of the Assembly Calculus involves five brain areas: areas *B* and *C* contain the assemblies *x, y* to be merged, while areas *B′* and *C′* contain the parents of these assemblies. The simultaneous spiking of the parents induces the firing of *x* and *y*, which then initiates a process whereby two-way afferent computation between *B* and *A*, as well as *C* and *A*, results in the adjustment of both *x* and *y* and the creation of an assembly *xy* in *A* having strong synaptic connections *to and from* both *x* and *y*. **G2:** Speed of convergence depends critically on *β*.

**Figure 2:**
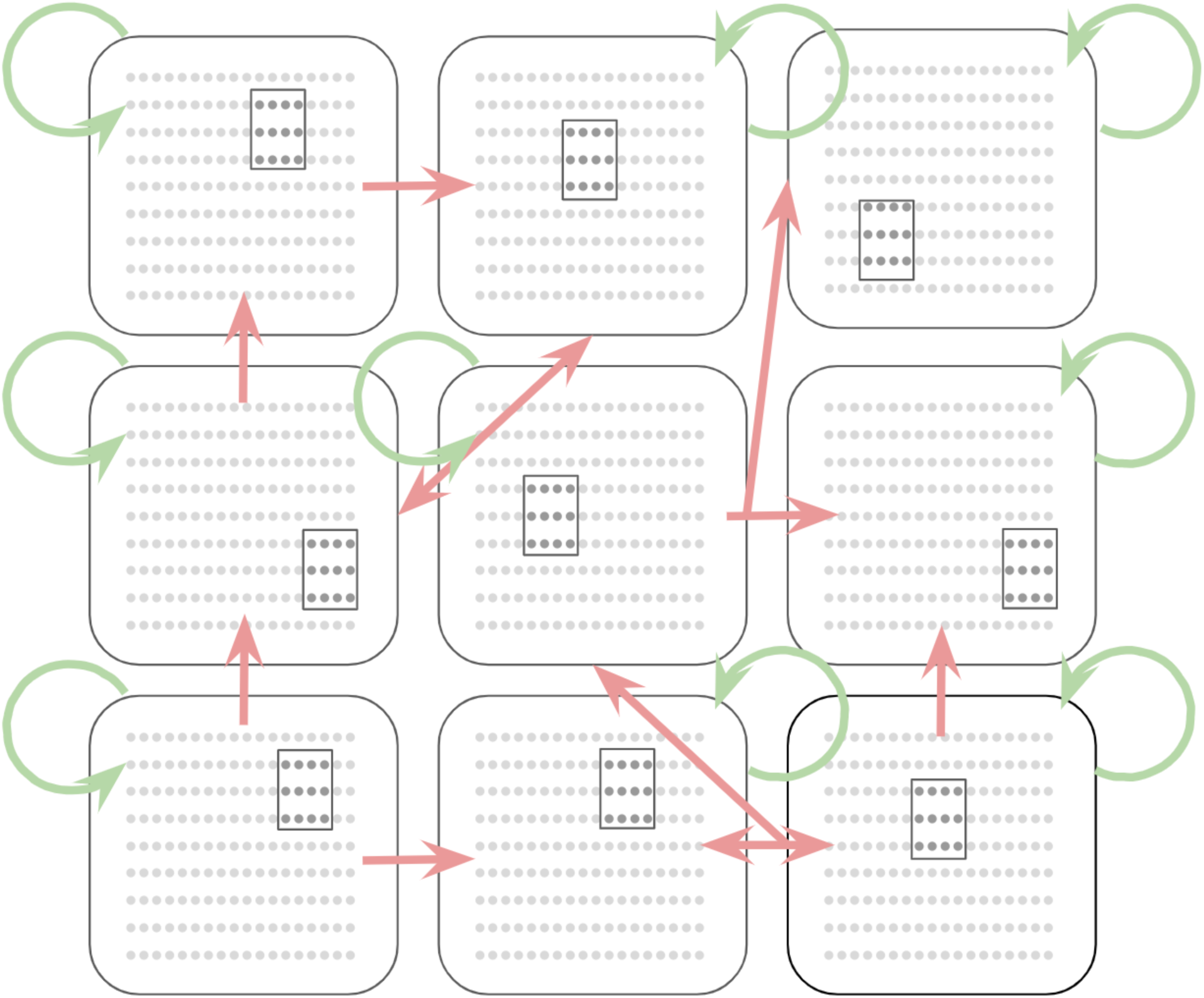
A mathematical model of the brain. Our analytical results, as well as most of our simulations, use a novel formal model of the brain, intended to capture cognitive phenomena in the association cortex. Our model encompasses a finite number of *brain areas* denoted *A, B*, … each containing *n* excitatory neurons. We assume that any pair of neurons in each area have a recurrent synaptic connection independently with probability *p*; and that for certain pairs of areas (*A, B*) there are also afferent connection from any neuron in *A* to any in *B*, also independently and with probability *p*. We assume that firing of neurons proceeds in discrete steps, and at each step *k* of the neurons in each area fire: namely, those *k* neurons which have the largest sum of presynaptic inputs from neurons that fired in the previous step. Synaptic weights are modified in a Hebbian fashion, in that synapses that fired (i.e., the presynaptic neuron fired and in the next step the postsynaptic neuron did) have their weight multiplied by (1 + *β*), where *β* is a plasticity constant. In our simulations we typically use *n* = 10^7^, *k* = 10^4^, *p* = 0.001, and *β* = 0.1.

We use this model to fathom quantitatively the creation and manipulation of assemblies of neurons. Since the model is probabilistic (by virtue of the random synaptic connectivity), our analytical results postulating the effectiveness of the various operations must contain the clause “with high probability,” where the event space is implicitly the underlying random graph. We assume that all cells in an assembly *x* belong to the same brain area, denoted `area`(*x*).

Our model also encompasses simplified forms of *plasticity and inhibition*. We assume multiplicative Hebbian plasticity: if at a time step neuron *i* fires and at the next time step neuron *j* fires, and there is a synapse from *i* to *j*, the weight of this synapse is multiplied by (1 + *β*), where *β >* 0 is the final parameter of our model (along with *n, p*, and *k*). Larger values of the plasticity coefficient *β* result in the operations converging faster, and render many of our proofs simpler. Finally, we model inhibition and excitatory–inhibitory balance by postulating that neurons fire in discrete time steps, and *at any time only a fixed number k of the n excitatory neurons in any area fire;* in particular, those neurons which receive the *k* highest excitatory inputs.

The four basic parameters of our model are these: *n* (the number of excitatory neurons in an area, and the basic parameter of our model); *p* (the probability of recurrent and afferent synaptic connectivity); *k* (the maximum number of firing neurons in any area); and the plasticity coefficient *β*. Typical values of these parameters in our simulations are *n* = 10^7^, *p* = 10^*-*3^, *k* = 10^4^, *β* = 0.1. We sometimes assume that *k* is (a small multiple of) the square root of *n*; this extra assumption seems compatible with experimental data, and yields certain interesting further insights.

Our model, as described so far, would result, through plasticity, in gigantic synaptic weights after a long time of operation. We further assume that synaptic weights are renormalized, at a slower time scale, so that the sum of presynaptic weights at each neuron stays relatively stable. This process of *homeostasis through renormalization* is orthogonal to the phenomena we describe here, and it interferes minimally with our arguments and simulations.

We emphasize that our model is *generic* in the sense that it is not assumed that circuits specific to various tasks are already in place. Its functionality — the needed apparatus for each task such as implementing an assembly operation — emerges from the randomness of the network and the selection of the *k* neurons with highest synaptic input as an almost certain consequence of certain simple events — such as the repeated firing of an assembly.

### Assembly projection

How do assemblies in the association cortex come about? It has been hypothesized (see e.g. [34]) that an assembly imprinting, for example, a familiar face in a subject’s medial temporal lobe (MTL) is created by the *projection* of a neuronal population, perhaps in the inferotemporal cortex (IT), encoding this face as a whole object. By *projection* of an assembly *x* we mean the creation of an assembly *y* in a downstream area that can be thought of as a “copy” of *x*, and such that *y* will henceforth fire every time *x* fires.

How is the new assembly *y* formed in a downstream area *B* by the repeated firing of *x* in area *A*? The process was vividly predicted in the discussion section of [14], for the case in which *A* is the olfactory bulb and *B* the piriform cortex. Once *x* has fired once, synaptic connectivity from area *A* to area *B* excites many neurons in area *B*. Inhibition will soon limit the firing in area *B* to a smaller set of neurons, let us call it *y*_1_, consisting in our framework of *k* neurons (see Figure 1A). Next, the simultaneous firing of *x* and *y*_1_ creates a stronger excitation in area *B* (one extra reason for this is *plasticity*, which has already strengthened the connections from *x* to *y*_1_), and as a result a new set of *k* neurons from area *B* will be selected to fire, call it *y*_2_. One would expect *y*_2_ to overlap substantially with *y*_1_ — this overlap can be calculated in our mathematical model to be roughly 50% for a broad range of parameters. If *x* continues firing, a sequence {*y*_*t*_} of sets of neurons of size *k* in area *B* will be created. For a large range of parameters and for high enough plasticity, this process can be proved to converge exponentially fast, with high probability, to create an assembly *y*, the result of the projection.

We denote the projection process described above as `project`(*x, B, y*). Assembly projection has been demonstrated both analytically [24, 29] and through detailed simulations [33, 26]; simulation results in our model, as well as improved analysis, are presented in Section 2, Figure 1.B4. Once the `project`(*x, B, y*) operation has taken place, we write *B* = `area`(*y*) and *x* = `parent`(*y*).

### Dense intraconnection of assemblies

Hebb [20] hypothesized that assemblies are *densely intraconnected* — that is, the chance that two neurons have a synaptic connection is significantly larger when they belong to the same assembly than when they do not — and our analysis and simulations verify this hypothesis (see Figure 1C). From the point of view of computation this is rather surprising, because the problem of finding a dense subgraph of a certain size in a sparse graph is a known difficult problem in computer science [12], and thus the very existence of an assembly may seem surprising. How can the brain deal with this difficult computational problem? The creation of an assembly through projection as outlined in the previous paragraphs provides an explanation: Since the elements of assembly *y* were selected to have strong synaptic input from the union of *x* and *y*, one intuitively expects the synaptic recurrent connectivity of *y* to be higher than random. In addition, the weights of these synapses should be higher than average because of plasticity.

### The RP&C primitive

It can be said that assembly operations, as described here, are powered exclusively by two forces known to be crucial for life more generally: *randomness* and *selection* (in addition to plasticity). No special-purpose neural circuits are required to be in place; all that is needed is *random synaptic connectivity* between, and recurrently within, areas; and selection, through inhibition in each area, of the *k* out of *n* cells currently receiving highest synaptic input. All assembly computation described here consists of applications of this operator, which we call *random projection and cap (RC&P)*. We believe that RP&C is an important primitive of neural computation, and computational learning more generally, and can be shown to have a number of potentially useful properties. For example, we establish analytically and through simulations that RP&C preserves overlap of assemblies remarkably well (as first noticed empirically in [7]). Finally, we used RP&C in experiments as the nonlinearity in each layer of a deep neural network, in the place of the sigmoid or the ReLU, and we found that it seems to perform at a level competitive with these.

### Association and pattern completion

In a recent experiment [21], electrocorticography (eCoG) recordings of human subjects revealed that a neuron in a subject’s MTL consistently responding to the image of a particular familiar place — such as the Pyramids — starts to also respond to the image of a particular familiar person — say, the subject’s sibling — once a combined image has been shown of this person in that place. A compelling parsimonious explanation of this and many similar results is that two assemblies imprinting two different entities adapt to the cooccurrence, or other observed affinity, of the entities they imprint by increasing their overlap, with cells from each migrating to the other while other cells leave the assemblies to maintain its size to *k*; we say that the two assemblies are *associating* with one another. The association of two assemblies *x* and *y in the same brain area* is denoted `associate`(*x, y*), with the common area `area`(*x*) = `area`(*y*) implicit. We can show analytically and through simulations that the simultaneous sustained firing of the two parents of *x* and *y* does effect such increase in overlap, while similar results have been obtained by simulations of networks of spiking neurons through STDP [33].

Assemblies are large and in many ways random sets of neurons, and as a result any two of them, if in the same area, may overlap a little by chance. If the assembly size *k* is about the square root of *n*, as we often assume in our simulations and quantitative analysis, this random overlap should be at most a very small number of neurons. In contrast, overlap resulting from the operation `associate`(*x, y*) is quite substantial: the results of [21] suggest an overlap between *associated* assemblies in the MTL of about 8 *-* 10% of the size of an assembly. The association between assemblies evokes a conception of a brain area as the arena of complex association patterns between the area’s assemblies; for a discussion of certain mathematical aspects of this view see [3].

One important and well studied phenomenon involving assemblies is *pattern completion:* the firing of the whole assembly *x* in response to the firing of a small subset of its cells [27]; presumably, such completion happens with a certain a priori probability depending on the particular subset firing. In our experiments pattern completion happens in a rather striking way, with small parts of the assembly being able to complete very accurately the whole assembly (see Figure 1.F1 and F2).

We believe that association and pattern completion open up fascinating possibilities for a genre of probabilistic computation through assemblies, a research direction which should be further pursued.

### Merge

The most sophisticated and complex operation in the repertoire of the Assembly Calculus is *merge*. Denoted `merge`(*x, y, A, z*), this operation starts with two assemblies *x* and *y*, in different brain areas, and creates a new assembly *z*, in a third area *A*, such that there is ample synaptic connectivity from *x* and *y* to *z*, and also vice versa, from *z* to both *x* and *y*.

Linguists had long predicted that the human brain is capable of combining, in a particularly strong sense, two separate entities to create a new entity representing this specific combination [4, 19], and that this ability is *recursive* in that the combined entity can in turn be combined with others. This is a crucial step in the creation of the hierarchies (trees of entities) that seem necessary for the syntactic processing of language, but also for hierarchical thinking more generally (e.g., deduction, discourse, planning, story-telling, etc.). Recent fMRI experiments [40] have indicated that, indeed, the completion of phrases and sentences (the completion of auditory stimuli such as “hill top” and “ship sunk”) activates parts of Broca’s area — in particular, the pars opercularis BA 44 for phrases, and the pars triangularis BA 45 for sentences. In contrast, unstructured word sequences such as “hill ship” do not seem to activate Broca’s area. Recall that Broca’s area has long been believed to be implicated in the syntactic processing of language.

A parsimonious explanation of these findings is that phrases and sentences are represented by assemblies in Broca’s area that are the results of the `merge` of assemblies representing their constituents (that is, assemblies for words such as “ship” and “sunk”); presumably the word assemblies reside in Wernicke’s area implicated in word selection in language. As these hierarchies need to be traversed both in the upward and in the downward direction (e.g., in the processes of language parsing and language generation, respectively), it is natural to assume that merge must have *two-way connections* between the new assembly and the constituent assemblies.

Our algorithm for implementing `merge,` explained in Section 2, is by far the most complex in this work, as it involves the coordination of *five* different brain areas with ample reciprocal connectivity between them, and requires stronger plasticity than other operations; our simulation results are reported in Section 2, see also Figure 1.G1 –G2.

Finally, a simpler operation with similar yet milder complexity is `reciprocal.project` (*x, A, y*): It is an extension of `project`(*x, A, y*), with the additional functionality that the resulting *y* has strong backward synaptic connectivity to *x*. Reciprocal projection has been hypothesized to be instrumental for implementing *variable binding* in the Brain — such as designating “cats” as the subject of the sentence “cats chase mice,” see [26]. The plausibility of `reciprocal.project` has been experimentally verified through detailed simulations of networks of spiking neurons with STDP [26], as well as in our simplified model.

### Readout and control operations

The purpose of the Assembly Calculus is to provide a formal system within which high level cognitive behavior can be expressed. Ultimately, we want to be able to write meaningful *programs* — in fact, parallel programs — in this system, for example containing segments such as:

~~~
if read(*A*) is null then project(*x, A, y*).
~~~

With this goal in mind, we next introduce certain additional low-level *control operations*, sensing and affecting the state of the system.

First, a simple *read* operation. In an influential review paper on assemblies, Buzsáki [5] proposes that, for assemblies to be functionally useful, readout mechanisms must exist that sense the current state of the assembly system and trigger appropriate further action. Accordingly, the Assembly Calculus contains an operation `read`(*A*) that identifies the assembly which has just fired in area *A*, and returns `null` othewise.

Finally, the Assembly Calculus contains two simple control operations. We assume that an assembly *x* in an area *A* can be explicitly caused to fire by the operation `fire`(*x*). That is to say, at the time an assembly *x* is created, a mechanism is set in place for igniting it; in view of the phenomenon of pattern completion discussed above, in which firing a tiny fragment of an assembly leads to the whole assembly firing, this does not seem implausible. We also assume that by default the excitatory cells in an area *A* are inhibited, unless explicitly disinhibited by the operation `disinhibit`(*A*) for a limited period of time, whose end is marked by the operation `inhibit`(*A*); the plausibility of the disinhibition–inhibition primitives is argued in [25] in terms of VIP cells [18].

#### Example

For a simple example of a program in the assembly calculus, the command `project`(*x, B, y*) where `area`(*x*) = *A*, is equivalent to the following:

~~~
disinhibit(*B*)
repeat *T*times: fire (*x*)
inhibit(*B*)
~~~

Simulations show that, with typical parameters, a stable assembly is formed after about *T* = 10 steps. An alternative version of this program, relying on the function `read`, is the following:

~~~
disinhibit(*B*)
repeat fire(*x*) until read(*B*) is not null
inhibit(*B*)
~~~

For another simple example, the command `associate`(*x, y*) whose effect is to increase the overlap of two assemblies in the same area *A* is tantamount to this:

~~~
disinhibit(*A*)
repeat *T* times: do in parallel {fire(parent((*x*)), fire(parent(*y*))}
inhibit(*A*)
~~~

A far more elaborate program in the Assembly Calculus, for a proposed implementation of the creation of *syntax trees* in the generation of natural language, is described in the discussion section.

### Computational power

It can be proved that the Assembly Calculus as defined above is capable of implementing, under appropriate assumptions, *arbitrary computations on O*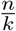 *bits of memory*. Given the intended values of *n* and *k*, this is a rather remarkable computational capability, suggesting — in view of the well established equivalence between parallel computation and space-bounded computation [28] — that any parallel computation with several hundred parallel steps, and on several hundred registers, can in principle be carried out by the Assembly Calculus.

### The assembly hypothesis

The Assembly Calculus is a formal system with a repertoire of rather sophisticated operations, where each of these operations can be ultimately reduced to the firing of randomly connected populations of excitatory neurons with inhibition and Hebbian plasticity. The ensuing computational power of the Assembly Calculus may embolden one to hypothesize that such a computational system — or rather a neural system far less structured and precise, which however can be usefully abstracted this way — underlies advanced cognitive functions, especially in the human brain, such as reasoning, planning, and language.

### Related work

Assemblies of excitatory neurons are, of course, not new: They have been hypothesized [20, 2], identified in vivo [17, 5], studied experimentally [27] and discussed extensively over the past decades — even, occasionally, in connection to computation, see [35]. However, we are not aware of previous work in which assemblies, with a suite of operations, are proposed as the basis of a computational system intended to explain cognitive phenomena.

Assemblies and their operations as treated here bear a certain resemblance to the research tradition of *hyperdimensional computing*, see [32, 22, 23], systems of high-dimensional sparse vectors equipped with algebraic operations, typically component-wise addition, component-wise multiplication, and permutation of dimensions. Indeed, an assembly can be thought as a high-dimensional vector, namely the characteristic vector of its support; but this is where the similarity ends. While assembly operations as introduced here are meant to model and predict cognitive function in the Brain, hyperdimensional computing is a machine learning framework — inspired of course by the Brain, like many other such frameworks — and used successfully, for example, for learning semantics in natural language [23]. In sparse vector systems there is no underlying synaptic connectivity between the dimensions, or partition of the dimensions into Brain areas. Finally, the operations of the Assembly calculus (project, associate, merge) are very different in nature, style, detail, and intent from the operations in sparse vector systems (add, multiply, permute).

Assembly computation is closer in spirit to Valiant’s model of neuroids and items [38], which was an important source of inspiration for this work. One difference is that whereas in Valiant’s model the neuron (called neuroid there) has considerable local computational power — for example, to set its pre-synaptic weights to arbitrary values —, in our formalism computational power comes from a minimalistic model of inhibition and plasticity; both models assume random connectivity. Another important difference is that, in contrast to an item in Valiant’s theory, an assembly is densely intraconnected, while the mechanism for its creation is described explicitly.

## 2 Results and Methods

### Projection

The operation `project`(*x, B, y*) entails activating repeatedly assembly *x* while *B* is disinhibited. Such repeated activation creates in the disinhibited area *B* a sequence of sets of *k* cells, let us call them *y*_1_, *y*_2_, …, *y*_*t*_, … The mathematical details are quite involved, but the intuition is the following: Cells in *B* can be thought of as competing for synaptic input. At the first step, only *x* provides synaptic input, and thus *y*_1_ consists of the *k* cells in *B* which happen to have the highest sum of synaptic weights originating in *x* — note that these weights are subsequently increased by a factor of (1 + *β*) due to plasticity. At the second step, neurons both in *x* and *y*_1_ spike, and as a result a new set *y*_2_ of “winners” from among cells of *B* is selected; for typical parameters, *y*_2_ overlaps heavily with *y*_1_. This continues as long as *x* keeps firing, with certain cells in *y*_*t*_ replaced by either “new winners” — cells that never participated in a *y*_*t′*_ with *t′ < t* — or by “old winners,” cells that did participate in some *y*_*t′*_ with *t′ < t*. We say that the process has *converged* when when there are no new winners. Upon further firing of *x, y*_*t*_ may evolve further slowly, or cycle periodically, with past winners coming in and out of *y*_*t*_; in fact, this mode of assembly firing (cells of the assembly alternating in firing) is very much in accordance with how assemblies have been observed to fire in Ca+ imaging experiments in mice, see for example [6]. It is theoretically possible that a new winner cell may come up after convergence; but it can be proved that this is a highly unlikely event, and we have never noticed it in our simulations. The number of steps required for convergence depends on the parameters, but most crucially on the plasticity coefficient *β*; this dependence is fathomed analytically and through simulations in Figure 1.B4).

### Density

It was postulated by Hebb [20] that assemblies are *densely interconnected* — presumably such density was thought to cause their synchronized firing. Since assemblies are created by projection, increased density is intuitively expected: cells in *y*_*t*_ are selected to be highly connected to *y*_*t-*1_, which is increasingly closer to *y*_*t*_ as *t* increases. It is also observed in our simulations (see Figure 1C) and predicted analytically in our model.

### Association and pattern completion

Our simulations (Figure 1E), as well as analytical results, show that the overlap of two assemblies *x, y* in the same area increases substantially in response of simultaneous firing of the two parent assemblies `parent`(*x*) and `parent`(*y*) (assumed to be in two different areas). The amount of post-association overlap observed in our simulations is compatible with the estimates in the literature [36, 8], and increases with the extent of co-occurrence (the number of consecutive simultaneous activations of the two parents.

Since association between assemblies is thought to capture affinity of various sorts, the question arises: If two associated assemblies *x* and *y* are both projected in another area, will the size of their overlap be preserved? Our results, both analytical and through simulations, strongly suggest that assembly overlap is indeed conserved reasonably well under projection (see Section 2 and Figure 1,). This is important, because it means that the signal of affinity between two assemblies is not lost when the two assemblies are further projected in a new brain area.

*Pattern completion* involves the firing, *with some probability*, of an assembly *x* following the firing of a few of its neurons [27, 6]. Simulations in our model (Figure 1.F2) show that, indeed, if the creation of an assembly is completed with a sufficient number of activations of its parent, then by firing fewer than half of the neurons of the assembly will result in most, or essentially all, of the assembly firing.

### Reciprocal projection and Merge

*Reciprocal projection*, denoted `reciprocal.project` (*x, B, y*), is a more elaborate version of projection. It involves `parent`(*x*) firing, which causes *x* in area *A* to fire, which in turn causes, as with ordinary projection, a set *y*_1_ in *B* to fire. The difference is that now there is synaptic connectivity from area *B* to area *A* (in addition to connectivity from *A* to *B*), which causes in the next step *x* to move slightly to a new assembly *x*_1_, while *y*_1_ has become *y*_2_. This continuing interaction between the *x*_*t*_’s and the *y*_*t±*1_’s eventually converges, albeit slower than with ordinary projection, and under conditions of ampler synaptic connectivity and plasticity. The resulting assembly *y* has strong synaptic connectivity both to and from *x* (instead of only from *x* to *y*, as is the case with ordinary projection). That reciprocal projection works as described above has been shown both analytically and through simulations in our model, as well as in simulations in a more realistic neural model with explicit inhibitory neurons and STDP in [25].

The operation `merge`(*x, y, A, z*) is essentially a double reciprocal projection. It involves the simultaneous repeated firing, in different areas, of the parents of both *x* and *y*, which causes the simultaneous repeated firing, also in two different areas *B* and *C*, of *x* and *y*. In the presence of enabled afferent two-way connectivity between *A* and *B*, and also between *A* and *C*, this initiates a process whereby a new assembly *z* is eventually formed in area *A*, which through its firing modifies the original assemblies *x* and *y*. In the resulting assemblies there is strong two-way synaptic connectivity between *x* and *z*, as well as between *y* and *z*. Analytical and simulation results are similar to those for reciprocal projection (see Figure 1.G1).

### Simulations

We gain insights into the workings of our model, and validate our analytical results, through simulations. In a typical simulation task, we need to simulate a number of discrete time steps in two or three areas, in a random graph with *∼ n*^2^*p* nodes (where the notation *∼ f* means”a small constant multiple of *f*). Creating this graph requires *∼ n*^2^*p* computation. Next, simulating each firing step entails selecting, in each area, the *k* out of *n* cells that receive that largest synaptic input. This takes ∼ *knp* computation per step. Since the number of steps needed for convergence are typically much smaller than the ratio 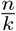, the *n*^2^*p* computation for the creation of the graph dominates the computational requirements of the whole simulation (recall that *n >> k*).

In our simulator we employ a maneuver which reduces this requirement to *∼ knp*, enabling simulations of the required scale on a laptop. The trick is this: We do not generate all *∼ n*^2^*p* edges of the random graph a priori, but generate them “on demand” as the simulation proceeds.

Once we know which cells fired at the previous step, we generate the *k* cells in the area of interest that receive the most synaptic input, as well as their incoming synapses from the firing cells, by sampling from the tail of the appropriate Bernoulli distribution; we then compare with previously generated cells in the same area to select the *k* cells that will fire next.

## 3 Discussion

We have defined a formal system intended to model the computations underlying cognitive functions. Its elementary object is an *assembly*, a set of excitatory neurons all belonging to the same brain area, and capable of near-simultaneous firing. The operations of this system enable an assembly to create a “copy” of itself in a new brain area through projection; two assemblies to increase their overlap in order to reflect observed co-occurrence in the world or other similarity; and, furthermore, two assemblies to create a new assembly with ample connectivity to and from the two original ones. These operations do not rely on pre-existing neural circuits; instead, their apparatus is generated on-the-fly, with high probability, through plasticity and randomness of synaptic connectivity. The resulting formal system, equipped with certain simple control operations, is fraught with considerable computational power. Central to our work is the speculation that something akin to this formal system may underly cognitive functions, especially in the human brain.

What is an assembly, exactly? The precise definition is a conundrum. In our narrative so far, assembly is simply a set of *k* excitatory neurons in a brain area capable of synchronous firing. According to Hebb’s prediction [20] as well as current observations [27] and neurorealistic simulations [33], the neurons of an assembly are not all activated simultaneously, but instead fire in a“pattern” with different neurons firing at different times^3^. In our formalism, and in our analytical proofs and our experiments, the assumption of discrete time steps suppresses to a large extent this sequential behavior. But even in our model, once the projection `project`(*x, B, y*) has stabilized (no new neurons in area *B* fire that had never fired before), a small number of neurons often keep coming in and out of the spiking set *y*_*t*_. One possible principled definition of an assembly is this: an assembly in an area *B* is a sparse distribution on the neurons of area *B*, whose support (the set of neurons with non-zero probability) is not much larger than *k*.

One novel aspect of our assembly operations is the crucial role of plasticity. While plasticity is well studied as an important aspect of the way the brain works and learning happens, we are not aware of many models in which plasticity takes place at the sub-second time scale, as it is hypothesized to do in assembly operations. Our use of assemblies as the basis of a computational system also departs from the usual discourse on assemblies, typically considered, implicitly, as fairly stable representations. In contrast, here we conjecture that assemblies can be formed and modified by the human brain at 4 Hz, the frequency of syllables in language (see the discussion on language below).

Our results and simulations assume uniformly random synaptic connectivity; however, experimental measurements [37, 16] suggest a departure from uniformity (but not from randomness). Our analytical results can be extended routinely to non-uniform random synaptic connectivity of this kind. In fact, our conclusions regarding the density and stability of assemblies can be further strengthened in such regimes. For example, suppose that, conditioned on the existence of two directed edges (*a, b*) and (*a, c*), the presence of edge (*b, c*) is much more likely, as concluded in [37]. Depending on the precise values of the parameters *n, k, p*, this would likely trigger a “birthday paradox” phenomenon (existence of two assembly cells *b, c* with synaptic connections from the same cell *a* of the parent assembly) that would further enhance the synaptic density, and hence the stability, of assemblies.

The basic operations of the Assembly Calculus as presented here — projection, association, reciprocal projection, and merge — correspond to neural population events which (1) are plausible, in the sense that they can be reproduced in simulations and predicted by mathematical analysis, and (2) provide parsimonious explanations of experimental results (for the merge and reciprocal project operations see the discussion of language below). In contrast, the read and control operations — read, fire, disinhibit —, however simple and elementary, lack in such justification, and were added for the purpose of rendering the Assembly Calculus a programmable computational system. Replacing them with more biologically plausible control operations leading to the same effect would be very interesting.

### Assemblies and Language

We hypothesized that assemblies and their operations may be involved in mediating higher cognitive functions in humans. In ending, we speculate below on how assemblies may be implicated in *language*.

Linguists have long argued that the language faculty is unique to humans, and that the human brain’s capability for syntax (i.e., for mastering the rules that govern the structure of sentences in a language) lies at the core of this ability [19, 4]. In particular, the linguistic primitive *Merge* has been proposed as a minimalistic basis of syntax. Merge combines two syntactic units to form one new syntactic unit; recursive applications of Merge can create a *syntax tree*, which captures the structure of a sentence (see Figure 3(a)). If this theory is correct, then which are the brain correlates of syntax and Merge? Recent experiments provide some clues — see [15] for an extremely comprehensive and informative recent treatment of the subject.

**Figure 3:**
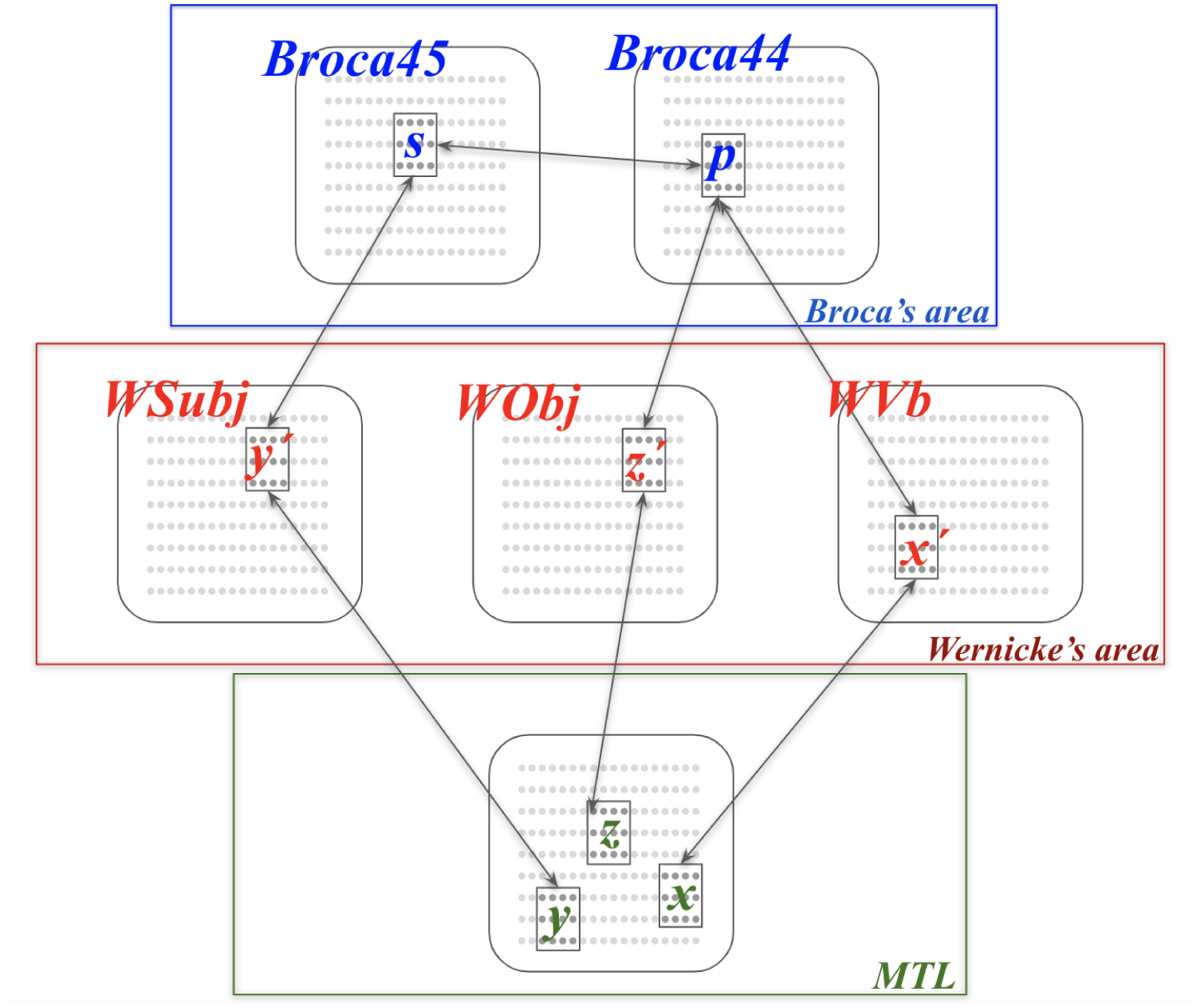
A proposed cortical architecture for syntax in language (see also [15]). In order to generate a simple sentence such as *“the boy kicks the ball,”* the subject must first *find* in the left MTL the representations of the verb, subject and object of the sentence. Next, these three assemblies are reciprocal-projected to the corresponding subareas of Wernicke’s area. Next the verb and the object are merged in BA 44 of Broca’s area to form a representation *p* of the verb phrase *“kicks the ball”*, and finally *p* is merged with the subject to create a representation *s* of the whole sentence in BA 45. This concludes the phase of building of the sentence’s *syntax tree*. Now in order to articulate the sentence, *s* fires and this results in the firing of the constituents of the tree, until eventually the three words in the MTL fire *in the correct order* for the subject’s language an order learned at infancy (in English, subject-verb-object). This latter activation of the three words mobilizes the corresponding motor functions resulting in the production of sound.

- Two different brain areas of the left superior temporal gyrus (which contains Wernicke’s area, known since the 19th century to be implicated in the use of words in language) seem to respond to the subject *vs* the object of a presented sentence [13];
- The completion of phrases and sentences presented is marked by activity in Broca’s area in the left hemisphere [39], known to be implicated in syntax;
- A sequence of sentences consisting of four monosyllabic words (such as “bad cats eat fish”) presented at the rate of four words per second (the natural frequency of articulated syllables) elicit a pattern of brain responses, with energy peaks at one, two, and four Hz, consistent with the hypothesis that syntax trees for the sentences are being constructed during the presentation [9].

If one accepts the hypothesis that indeed something akin to syntax trees is constructed in our brain during the parsing — and presumably also during the generation — of sentences, one must next ask: how is this accomplished? According to [15] Chapter 4, these and a plethora of other experimental results point to a *functional cortical network* for lexical and syntactic processing, involving the MTL, Broca’s areas BA 44 an BA 45, and Wernicke’s area in the superior temporal gyrus, as well as axon fibers connecting these four areas, all in the left hemisphere. Syntactic processing of language seems to entail a complex sequence of activations of these areas and transmission through these fibers. How is this orchestrated sequence of events carried out? Does it involve the generation and processing, on-the-fly, of representations of the constituents of language (words, phrases, sentences)? Angela Friederici [15] proposes in page 134 that “for language there are what I call *mirror neural ensembles* through which two distinct brain regions communicate with each other.” Could it be that assemblies and their operations play this role?

We propose a dynamic brain architecture for the generation of a simple sentence, powered by assemblies and the operations `reciprocal.project` and `merge`, and consistent with the experimental results and their interpretations outlined above (see Figure 3). In particular, we propose that the construction of the syntax tree of a simple sentence being generated by a subject can be implemented by the following program of the Assembly Calculus:

~~~
do in parallel: find-verb(*Im*, MTL, *x*), find-subj(*Im*, MTL, *y*), find-obj(*Im*, MTL, *z*);
do in parallel: reciprocal.project (*x*, WVb, *x′*),
    reciprocal.project (*y*, WSubj, *y′*), reciprocal.project (*x*, WObj, *z′*);
merge(*x′, z′*, Broca44, *p*);
merge(*y′, p*, Broca45, *s*)
~~~

The generation of a sentence such as *“the boy kicks the ball”* starts with a desire by the subject to assemble — and possibly articulate — a particular fact. The raw fact to be put together is denoted here by *Im* — an image, sensed or mental. In the first line, the subject’s lexicon, presumably a collection of tens of thousands of assemblies in the medial temporal lobe (MTL), is searched in order to identify the verb (the action in the fact relayed in *Im*); the subject (the agent of the fact); and the object (the patient of the fact). We assume that the corresponding brain functions `find-verb` etc. are already in place. For the example *“the boy kicks the ball”*, the three assemblies *x, y*, and *z* are identified, encoding the words *“kick”, “boy”*, and *“ball”*, respectively. In the second line, these three assemblies are projected to the three sub-areas of Wernicke’s area specializing in the verb, subject, and object of sentences, respectively. In fact, instead of ordinary projection, the `reciprocal.project` operation is used, as first proposed in [25], for reasons that will become clear soon. Next, an assembly *p* is formed in the pars opercularis of Broca’s area (BA 44) representing the verb phrase *“kicks the ball”* through the `merge` operation applied to the two assemblies *x′* and *z′* encoding the constituent words of the phrase in Wernicke’s area. Finally, an assembly *s* corresponding to the whole sentence is formed in the pars triangularis of Broca’s area (BA 45) via the `merge` of assemblies *p* and *y′*, completing the construction of the syntax tree of the whole sentence.

The Assembly Calculus program above accounts only for the first phase of sentence production, during which the syntax tree of the sentence is constructed. Next the sentence formed may be articulated, and we can speculate on how this process is carried out: Assembly *s* is activated, and this eventually causes the assemblies *x′, y′, z′* — the leaves of the syntax tree — to fire. The activation of these three assemblies is done in the order specific to the speaker’s language learned at infancy (for example, in English *subject-verb-object*); note that the first phase of building the syntax tree was largely language independent. Eventually, the lexicon assemblies will be activated in the correct order (this activation was the purpose of utilizing the `reciprocal.project` operation), and these in turn will activate the appropriate motor functions which will ultimately translate the word sequence into sounds.

The above narrative is only about the building of the basic syntactic structure — the “scaffold” of extremely simple sentences, and does not account for many other facets: how the three `find-` tasks in the first line are implemented; how the noun phrases are adorned by determiners such as “the” and how the verb is modified to reflect person or tense (“kicks” or “kicked”); and the important inverse processes of *parsing and comprehension*, among a myriad of other aspects of language that remain a mystery.

## Acknowledgments

The authors are grateful to Larry Abbott for his insightful ideas and comments concerning this work. We also wish to thank several colleagues for helpful discussions and feedback, including Richard Axel, Bob Berwick, Costis Daskalakis, Boris Harizanov, Pentti Kanerva, Robert Legenstein, Chris Manning, Bruno Olshausen, Tommy Poggio, Les Valiant, and Rafa Yuste. CHP’s research was partially supported by NSF awards CCF1763970 and CCF1910700 and by a research contract with Softbank; SSV’s by NSF awards 1717349, 1839323 and 1909756; and WM’s by the Human Brain Project grant 785907 of the European Union.

## Footnotes

1 Our assumption that all areas contain the same number of neurons is one of expository, not mathematical, convenience, and provides the basic numerical parameter *n* of our analysis. A slightly more complex version would have parameters *n*_*A*_, and *k*_*A*_ for each area *A*.

2 Several experimental results [37, 16] suggest deviations of the synaptic connectivity of the animal brain from the uniformity of *G*_*n,p*_. We discuss how such deviations affect — mostly support — our framework in the discussion section.

3 In musical metaphor, an assembly is thought to be not an octave but a cord sequence.

